# DDK has a primary role in processing stalled replication forks to initiate downstream checkpoint signaling

**DOI:** 10.1101/207266

**Authors:** Nanda Kumar Sasi, Flavie Coquel, Yea-Lih Lin, Jeffrey P MacKeigan, Philippe Pasero, Michael Weinreich

**Author notes:** To whom correspondence should be addressed: Michael Weinreich, PhD, Van Andel Research Institute, 333 Bostwick Ave. NE, Grand Rapids, MI 49503, USA.

## Abstract

CDC7-DBF4 kinase (DDK) is required to initiate DNA replication in eukaryotes by activating the replicative MCM helicase. DDK has also been reported to have diverse and sometimes conflicting roles in the replication checkpoint response in various organisms but the underlying mechanisms are far from settled. Here we show that human DDK promotes limited resection of newly synthesized DNA at stalled replication forks or sites of DNA damage to initiate replication checkpoint signaling. DDK is also required for efficient fork restart and G2/M cell cycle arrest. DDK exhibits genetic interactions with the ssDNA exonuclease EXO1, and we show that EXO1 is also required for nascent strand degradation following exposure to HU, raising the possibility that DDK regulates EXO1 directly. Thus, DDK has a primary and previously undescribed role in the replication checkpoint to promote ssDNA accumulation at stalled forks, which is required to initiate a robust checkpoint response and cell cycle arrest to maintain genome integrity.

---

DDK is a two-subunit kinase that is essential to initiate DNA replication at individual replication origins by phosphorylating and activating the MCM2-7 replicative helicase. The regulatory subunit DBF4 binds to CDC7 and is required for its kinase activity [1]. DDK also has roles in replication checkpoint signaling that are less well understood [1]. In budding yeast, DDK is a target of the checkpoint effector kinase Rad53 that is activated following replication stress [2]. Rad53-mediated phosphorylation of Dbf4 modestly reduces DDK activity [2] and also inhibits late origin firing [3, 4], but there is also evidence that DDK is required for the complete activation of Rad53 kinase [5]. In fission yeast, DDK subunits Hsk1 and Dfp1 are phosphorylated upon HU treatment by the checkpoint kinase Cds1, orthologous to budding yeast Rad53 and mammalian CHK2 [6–8]. Deletion of either Cds1 or its activating protein Mrc1 partially rescues the temperature sensitivity of *hsk1-1312* mutants suggesting that Cds1 (like Rad53) inhibits DDK activity [8, 9]. Cds1 activation in response to HU, however, was reduced significantly in *hsk1*(ts) strains at restrictive temperature and the cell cycle arrest was also aberrant as seen by an increased population of ‘cut’ cells (indicative of mitosis without complete DNA replication) [8, 10]. These paradoxical results in yeast could be explained by a negative feedback loop where DDK first helps to initiate replication checkpoint activation and then the checkpoint pathway subsequently alters DDK activity to inhibit late origin firing.

Initial studies in human cells showed that the replication checkpoint inhibits DDK activity, presumably to inhibit origin firing [11, 12]. However, subsequent studies reported that human DDK is active during replication stress and has an upstream role to fully activate the checkpoint kinase CHK1 [13, 14]. In response to exogenous replication inhibitors, DDK recruitment to chromatin is increased [13, 14], CDC7-DBF4 complex is stabilized [13], and MCM shows increased phosphorylation at several DDK-specific sites [13, 15]. DDK promotes CHK1 phosphorylation partly through binding to and phosphorylating Claspin, an adaptor protein required for CHK1 activation [16, 17]. Although depleting the CDC7 protein subunit using a CDC7 siRNA resulted in loss of chromatin-bound Claspin following HU treatment [16], inhibiting DDK activity using small molecule inhibitors did not [17]. Furthermore, depleting Claspin using siRNA did not reduce CHK1 phosphorylation to the same extent as CDC7 siRNA [16], suggesting that DDK could promote checkpoint signaling by other mechanisms. In this study, we show that DDK has a primary role to initiate checkpoint signaling by promoting ssDNA formation at stalled forks, most likely by promoting nascent DNA resection.

## Results and Discussion

### DDK has a primary role in initiating replication checkpoint signaling

Since blocking DDK activity largely prevents CHK1 phosphorylation upon exposure to replication stress [14, 16, 17], we analyzed how DDK influences the overall checkpoint response induced by HU. We found that 5 μM of DDKi and a pre-treatment time of 4 hours were required to substantially block CHK1 activation by HU in HCC1954 cells **(Supplementary Figure 1A)**. HCC1954 is a breast cancer cell line that expresses high levels of both DDK subunits and exhibits robust decrease in MCM2 phosphorylation in response to the prototype DDK inhibitor, PHA-767491 (DDKi) [18]. As shown previously, CHK1 activation was almost completely eliminated by pre-treatment with the DDK inhibitor PHA-767491 **(Figure 1A)**. A similar effect on CHK1 activation was seen in response to the more specific DDK inhibitor XL413 in Colo-205 cells **(Figure 1B)**. However unexpectedly, we found that RPA2 phosphorylation and chromatin accumulation following HU exposure were also significantly reduced if DDK activity was blocked **(Figure 1A, B)**. This is not consistent with a singular role for DDK to promote CHK1 activation through Claspin phosphorylation. We saw an identical decrease in RPA2 phosphorylation and chromatin accumulation when DDK activity was blocked using an siRNA against *CDC7* **(Supplementary Figure 1B).** Chromatin-bound RPA2 immunofluorescence, which is sharply increased after 2 hours of HU exposure, was also significantly reduced in CDC7 siRNA-treated cells **(Supplementary Figure 1C)**. DDK inhibition also prevented CHK1 phosphorylation, RPA2 accumulation and RPA2 phosphorylation in response to two DNA damaging agents, camptothecin and etoposide **(Supplementary Figure 1D, E)**. Using a non-denaturing BrdU assay that directly measures ssDNA formation we found that the ssDNA formation upon HU treatment was significantly reduced upon DDK inhibition **(Figure 1C)**. DDK inhibition alone, in the absence of exogenous replication inhibitors, did not result in widespread ssDNA formation nor the activation of replication-checkpoint signaling **(Supplementary Figure 2A, B)** confirming several earlier findings [19, 20]. RPA2 S33 was mildly phosphorylated in response to the DDK inhibitor as opposed to hyperphosphorylation observed upon replication stress [21]. Taken together, our results show that DDK inhibition substantially blocks the accumulation of ssDNA and downstream events (i.e., RPA binding to ssDNA) in response to replication fork stalling by HU or DNA damage by camptothecin and etoposide.

**Figure 1:**
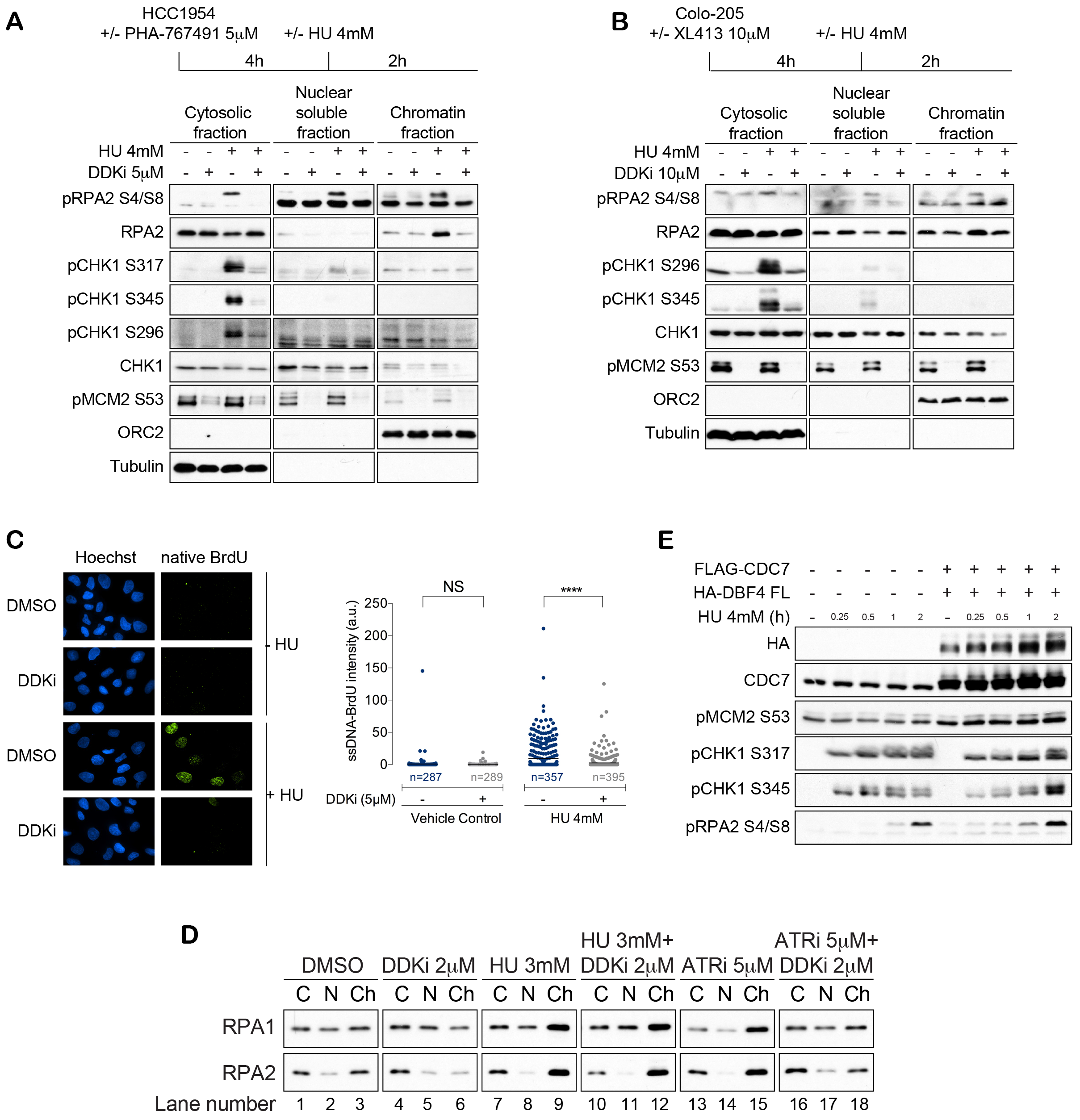
DDK activity is required for generating ssDNA and initiating the checkpoint pathway. **(A)** HCC1954 cells or **(B)** Colo-205 cells were pre-treated with DMSO or DDKi for 4h followed by incubation with or without HU for 2h, cell fractionation and immunoblot analysis. **(C)** BrdU-treated HCC1954 cells were treated as in (A) and analyzed using immunofluorescence microscopy. **(D)** HCC1954 cells were treated with indicated drugs for 2h followed by cell fractionation and immunoblot analysis. C - Cytoplasmic, N - Nuclear soluble, Ch - Chromatin fraction. **(E)** HEK-293T cells were transfected with indicated constructs and 48 hours later treated with HU for the indicated time.

While reduced origin firing upon DDK inhibition might explain the lack of checkpoint activation in response to HU, several pieces of evidence argue against this and suggest a more direct role for DDK at stalled forks. Firstly, inhibition of ATR kinase in unperturbed cells induces aberrant origin firing leading to accumulation of chromatin-bound RPA1 and RPA2 (**Figure 1D**, **lane 3 vs. 15**) [22, 23]. This RPA accumulation, however, was prevented upon co-treatment with DDKi (**Figure 1D**, **lane 15 vs. 18)** consistent with the essential role of DDK in origin firing. In contrast, HU-mediated RPA accumulation on chromatin, which is blocked by prior treatment with DDKi, was not affected by co-treatment with DDKi (**Figure 1D**, **lane 9 vs. 12)**. In fission yeast *hsk1-89* temperature sensitive strain that alters the Cdc7 subunit showed ~20-fold reduction in Cds1 (Rad53/CHK2) activation following HU treatment, even at permissive temperature [10]. Other fission yeast mutants defective in origin firing (e.g., *mcm4* or *orc1* mutants) did not show a similar reduction in Cds1 activation [10], suggesting an origin-independent role for Hsk1 (CDC7) in Cds1 activation. Budding yeast *cdc7* and *dbf4* mutants also have substantially reduced levels of Rad53 [24]. These results suggest that DDKi-mediated reduction in origin firing per se does not significantly affect the extent of checkpoint activation and ssDNA at stalled forks.

We also found that overexpressing human DDK subunits increased the phosphorylation of CHK1 and RPA2 in response to HU treatment **(Figure 1E)**, again consistent with a positive role for DDK in promoting checkpoint activation. In summary, our results indicate that DDK promotes formation of ssDNA-RPA complexes upon exposure to various forms of replication stress independently of its role in origin activation. This was confirmed below using single molecule analyses.

### DDK is required for processing stalled replication forks

Based on the findings described above, we hypothesized that DDK is required to process stalled replication forks to activate the replication checkpoint. To test our hypothesis, we performed DNA fiber analysis and measured the effect of inhibiting DDK at established forks with or without HU treatment **(Figure 2A)**. A shorter incorporation time (consecutive 20 mins pulses of IdU and CldU) allowed detection of smaller changes in the length of nascent DNA than in previously published assays (P. Pasero, unpublished data). Treatment with DDKi alone had no effect on CldU track length indicating that DDK inhibition does not alter nascent strand length in unperturbed cells (**Figure 2A** **and B, DMSO vs. DDKi 2h, DDKi 4h; Supplementary Figure 3A)**. Moreover, DDKi-treated samples did not show an appreciable decrease in the density of replication fibers (not shown) indicating that sufficient replication forks exist in these cells to activate a checkpoint response. We found that 2 hours of HU exposure significantly reduced the length of nascent DNA tracks (CldU tracks) in HCC1954 cells compared to untreated cells **(Supplementary Figure 3A, DMSO vs. DMSO+HU)**. A reduction in the ratio of CldU to IdU lengths was also observed upon 2 hours of HU treatment, which normalizes for any change in the rates of replication elongation (**Figure 2B**, **DMSO vs. DMSO+HU)**. Similar shortening of nascent DNA tracts was observed in HU-treated MCF-7 cells (**Supplementary Figure 3B-D)** and in several different cell lines irrespective of their genetic backgrounds, with tract-length shortening directly correlated with the time of incubation in HU (P. Pasero, unpublished data). Short exposure to HU does not induce cell death in HCC1954 cells **(Figure 4B)** and several studies have shown that collapse of stalled forks only occurs after prolonged (12 to 24hr) exposure to HU [27, 28]. Therefore, nascent strand degradation seen after 2h of HU exposure is likely to be a physiological response.

**Figure 2:**
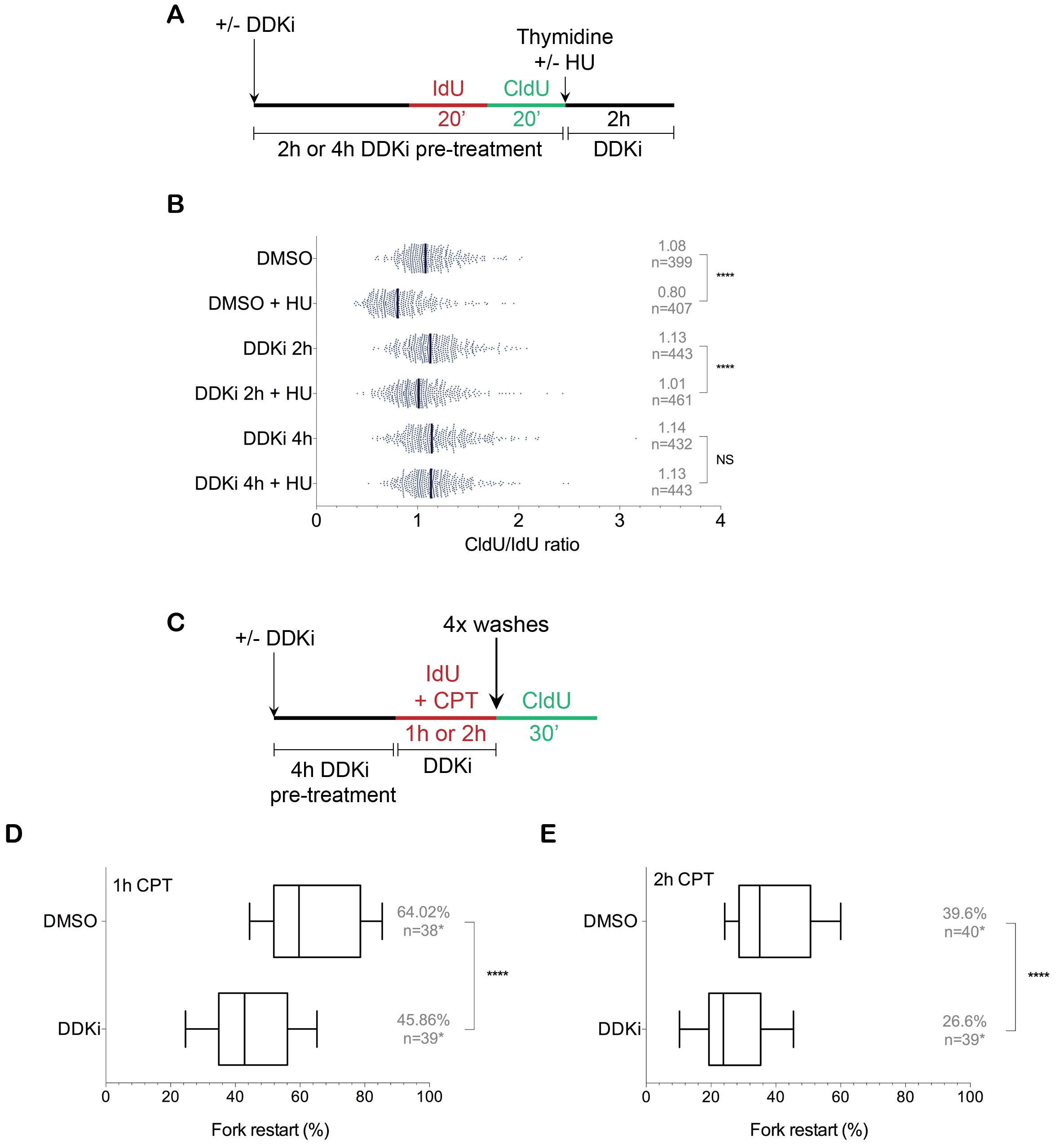
DDK has a primary role in processing and restarting stalled replication forks. **(A)** HCC1954 cells were pre-treated with DMSO or DDKi for 2h or 4h, labeled consecutively with IdU and CldU (20 mins each), subjected to a thymidine chase with or without HU for 2h (still in the presence of DDKi (shown) or DMSO), and then harvested for DNA fiber assay. **(B)** Nascent strand resection was measured as a ratio of CldU to IdU incorporation. **(C)** HCC1954 cells were pretreated with or without DDKi for 4h, exposed to camptothecin (CPT) for 1h or 2h in the presence of IdU, washed four times with PBS and exposed to CldU for a further 30 minutes. Replication fork restart was measured by counting DNA fibers with contiguous IdU and CldU tracks after exposure to 1h **(D)** or 2h **(E)** of CPT. n=xx* in restart assay indicates the number of images counted per sample. 25 to 30 fibers were counted per image.

Surprisingly, DDKi treatment prevented degradation of nascent DNA upon HU exposure (**Figure 2B**, **DMSO+HU vs. DDKi 2h/4h+HU; Supplementary Figure 3A)**. The reduction in nascent strand degradation was dependent on the extent of DDK inhibition with 4 hours of DDKi pre-treatment resulting in CldU track lengths similar to DMSO treated cells **(Figure 2B)**. We find that RPA accumulation on chromatin correlates well with the length of nascent DNA seen in DNA fiber assay **(Figure 1A, B and Figure 2B)**. Based on this data, we propose that DDK actively promotes limited resection of stalled replication forks, which is required for formation of ssDNA, local accumulation of RPA, and activation of downstream checkpoint signaling.

In a Xenopus *in vitro* system, aphidicolin treatment, which inhibits DNA polymerase alpha and delta, results in unrestricted DNA unwinding and ssDNA formation ahead of the stalled fork that is thought to induce the replication checkpoint [29]. This was proposed to be due to a functional (but not physical) uncoupling of the replicative polymerases and helicase **(see model in** **Figure 5**). Given that helicase and polymerase activities are strongly interdependent and depend on physical interactions in many model organisms [30–32], it is not clear how a functional uncoupling might result in independent helicase activity in living cells. Electron microscopic analysis has shown that ssDNA formed in yeast at stalled forks is limited to one parental strand [33, 34]. However, uncoupling between the polymerase and helicase would generate long ssDNA on both leading and lagging strands. In addition, diverse types of replication inhibitors generate ssDNA, even inter-strand crosslinking agents, which would not allow helicase activity or polymerase activity beyond the crosslink [35]. So, while helicase-polymerase uncoupling can generate ssDNA ahead of the forks in response to certain types of replication stress, our data suggest that DDK-dependent limited processing of nascent DNA *behind* stalled forks might be a more universal mechanism for ssDNA formation.

### DDK is required for restart of stalled replication forks

An important function of the replication checkpoint pathway is to promote DNA repair mechanisms required for rescuing and restarting stalled forks [1]. Using a DNA fiber assay, we measured the rate of fork restart following 1 or 2 hours of CPT exposure **(Figure 2C)**. If DDK is required for fork restart, we would expect to see a reduced rate of fork restart in CPT+DDKi treated cells compared to CPT treatment alone. Sixty-four percent of forks restarted after 1h of 1 μM CPT exposure and this was reduced to 46% when cells were pre-treated with DDKi for 4 hours **(Figure 2D)**. The effect of 2 hours of CPT on fork restart was more severe with only 40% of forks restarting after removal of CPT. This was further reduced to 27% when they were pre-treated with DDKi for 4 hours **(Figure 2E)**. These results confirm that DDK is required for efficient replication fork restart, presumably because cells are defective in initiating the replication checkpoint and DNA repair and therefore cannot efficiently restart stalled replication forks.

### DDK likely promotes fork processing by regulating the activity of nucleases at stalled forks

To identify potential nucleases and helicases required for processing replication forks after 2 hours exposure to HU, we used siRNAs to knock down enzymes known to act on collapsed forks plus those identified at unperturbed and stalled replication forks through iPOND analysis [36–38]. Knockdown of EXO1, BLM, and CtIP reduced phospho-CHK1 levels in response to HU whereas MRE11 knockdown did not (**Figure 3A** **and Supplementary Figure 4A)**. Knockdown of these enzymes did not significantly affect cell cycle profiles **(Supplementary Figure 4B)**. EXO1 knockdown showed the strongest effect on CHK1 phosphorylation. Moreover, in HEK-293 cells EXO1 knockdown significantly reduced ssDNA formation after 2h of HU treatment [39]. We did not observe a reduction in RPA2-S4/S8 phosphorylation upon knockdown of any of the potential resection enzymes. It has been suggested that the initial resection at stalled forks is independent of regression of nascent DNA at stalled forks, which could be the signal for RPA2-S4/S8 phosphorylation [28]. Consistent with this, knockdown of the fork regression helicase FBH1 resulted in a decrease in RPA2-S4/S8 phosphorylation after a short HU treatment but had no effect on ATR-CHK1 signaling [28]. Since DDK inhibition results in significant reduction in both CHK1 and RPA2 phosphorylation, this further argues for an upstream role for DDK in replication checkpoint activation.

EXO1 exists in a complex with EEPD1, BLM, and RPA and knockdown of individual proteins destabilizes other proteins in the complex [40]. Interestingly, EXO1 and BLM are significantly less abundant following exposure to DDKi with or without exposure to DNA damaging agents (**Figure 3B**, **bottom panels)**. Moreover, knocking down either CDC7 or EXO1 significantly reduced expression of the other protein **(Figure 3C)**. Destabilization of EXO1 could be rescued by treatment with the proteasome inhibitor MG-132, indicating that EXO1 is actively degraded post-transcriptionally in the absence of DDK activity **(Figure 3D)**. In contrast, CtIP stability was only slightly reduced upon DDK inhibition and the levels of MRE11 were unchanged **(Figure 3B)**. This data shows that DDK is required for EXO1 and BLM stability and therefore activity, and further suggests an important role for the EXO1-BLM complex in fork resection immediately after fork stalling.

**Figure 3:**
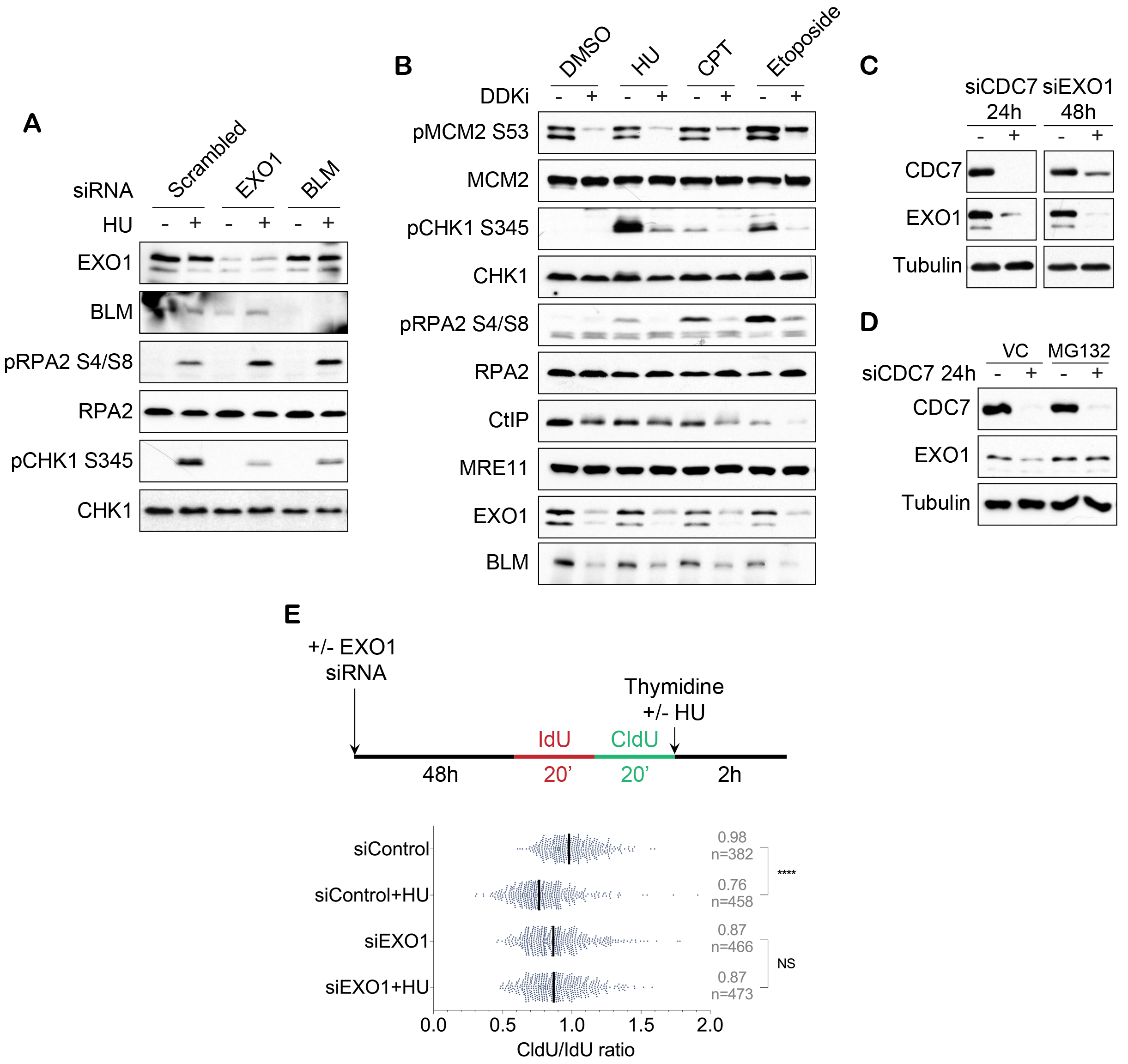
DDK could promote fork resection by directly regulating the activity of nucleases. **(A)** HCC1954 cells were transfected with indicated siRNAs, 48h later treated with or without HU for 2h. **(B)** HCC1954 cells were pre-treated with DMSO (-) or DDKi (+) for 4h followed by incubation with DMSO, HU, CPT or Etoposide for a further 2h. **(C)** HCC1954 cells were transfected with indicated siRNA for 24h or 48h. **(D)** HCC1954 cells were transfected with CDC7 siRNA, 24h later treated with a vehicle control (VC) or MG132 for 3h. All samples were analyzed by immunoblot analysis. **(E)** MCF-7 cells were transfected with control or EXO1 siRNA and 48 hours later subjected to DNA fiber assay as in 2A (CldU/IdU ratio).

To more directly assess the role of EXO1 in regulating nascent strand length when forks stall, we performed DNA fiber analysis in MCF-7 cells transfected with control or EXO1 siRNAs and measured nascent tract lengths after exposure to HU. While control siRNA cells showed the expected reduction in CldU/IdU ratio upon HU exposure, EXO1-depleted cells did not show a similar reduction indicating that nascent strand resection is reduced upon EXO1 knockdown **(Figure 3E)**. The cause of reduced basal CldU/IdU ratio in EXO1-depleted cells **(Figure 3E,** 0.87 in siEXO1 vs 0.98 in siControl) is not known. In a separate DNA fiber assay performed in the absence of thymidine chase, nascent DNA tracks were again unaffected by HU in EXO1-depleted cells **(Supplementary Figure 5A, B)** further implicating this nuclease in the nascent strand degradation following HU exposure. Very recently, Exo1 deletion in fission yeast was shown to prevent ssDNA and RPA accumulation at arrested replication forks [41] further substantiating our finding of a role for EXO1 in resection immediately after fork stalling in mammalian cells. We suggest that DDK recruitment to stalled forks facilitates limited nucleolytic removal of nascent DNA involving EXO1, which in turn generates ssDNA-RPA complexes and activates replication checkpoint signaling.

### Low dose DDKi causes aberrant mitotic structures

While untransformed human cells prevent S-phase progression upon DDK inhibition by inducing a G1/S-checkpoint [19, 42], tumor cells proceed through an aberrant S-phase and undergo apoptosis but the exact mechanism that triggers apoptosis remains unknown [19, 20]. Inhibiting DDK during S-phase will reduce the number of overall replication forks and inhibit restart of stalled forks (shown here) leading to incomplete DNA replication and a G2/M arrest. However, DDKi-treated cells are also significantly defective in checkpoint activation and therefore would have difficulty restraining mitosis. Our findings therefore prompted us to examine mitotic progression in HCC1954 cells using a low dose of DDKi that will allow S-phase progression. Strikingly, we found numerous aberrant mitotic figures in the DDKi-treated cells similar to cells treated with a low dose (0.4μM) of aphidicolin, which is known to slow DNA polymerization and induce mitotic abnormalities [43, 44] **(Figure 4A)**. The defects in mitosis observed after 24-hour treatment with 1μM DDKi (PHA) were not a by-product of apoptosis as the same effect was seen both with and without co-treatment with the pan-caspase inhibitor zVAD. To confirm that this effect is DDK-specific we also tested the more selective biochemical DDK inhibitor, XL413 [45]. In HCC1954 cells and many other cell lines, XL413 is a poor in vivo inhibitor of DDK activity and has little effect on cell growth even at high inhibitor concentrations [18]. We therefore used relatively high 10μM and 20μM concentrations of XL413 to moderately inhibit DDK in HCC1954 cells but not induce apoptosis. We found that XL413 treatment also significantly increased the number of mitotic abnormalities in a dose-dependent manner confirming this to be a DDK-specific phenotype **(Figure 4A)**. Although the increase in lagging chromosomes was similar in aphidicolin and DDKi-treated cells, DDKi treatment resulted in higher number of anaphase bridges compared to aphidicolin-treated cells. Anaphase bridges are thought to arise from chromosomes that have long stretches of incompletely replicated DNA, fused telomeres, or chromatid cohesion defects [46]. Our data suggest that low-level DDKi-treated cells undergo mitosis in presence of under-replicated DNA. Correspondingly, cell cycle arrest by HU pretreatment rescued cell death induced by DDKi and the extent of rescue was positively correlated with increasing time in HU, with 24 hours of HU pre-treatment showing the strongest rescue **(Figure 4B, C)**. Inhibition of CHK1 activation using a specific CHK1 inhibitor (LY2603618) did not abrogate the rescue seen upon HU pre-treatment **(Figure 4D, E)**. Therefore, preventing S-phase or fork progression protects cells against apoptosis upon DDK inhibition, but active CHK1 is not sufficient for this protection.

**Figure 4:**
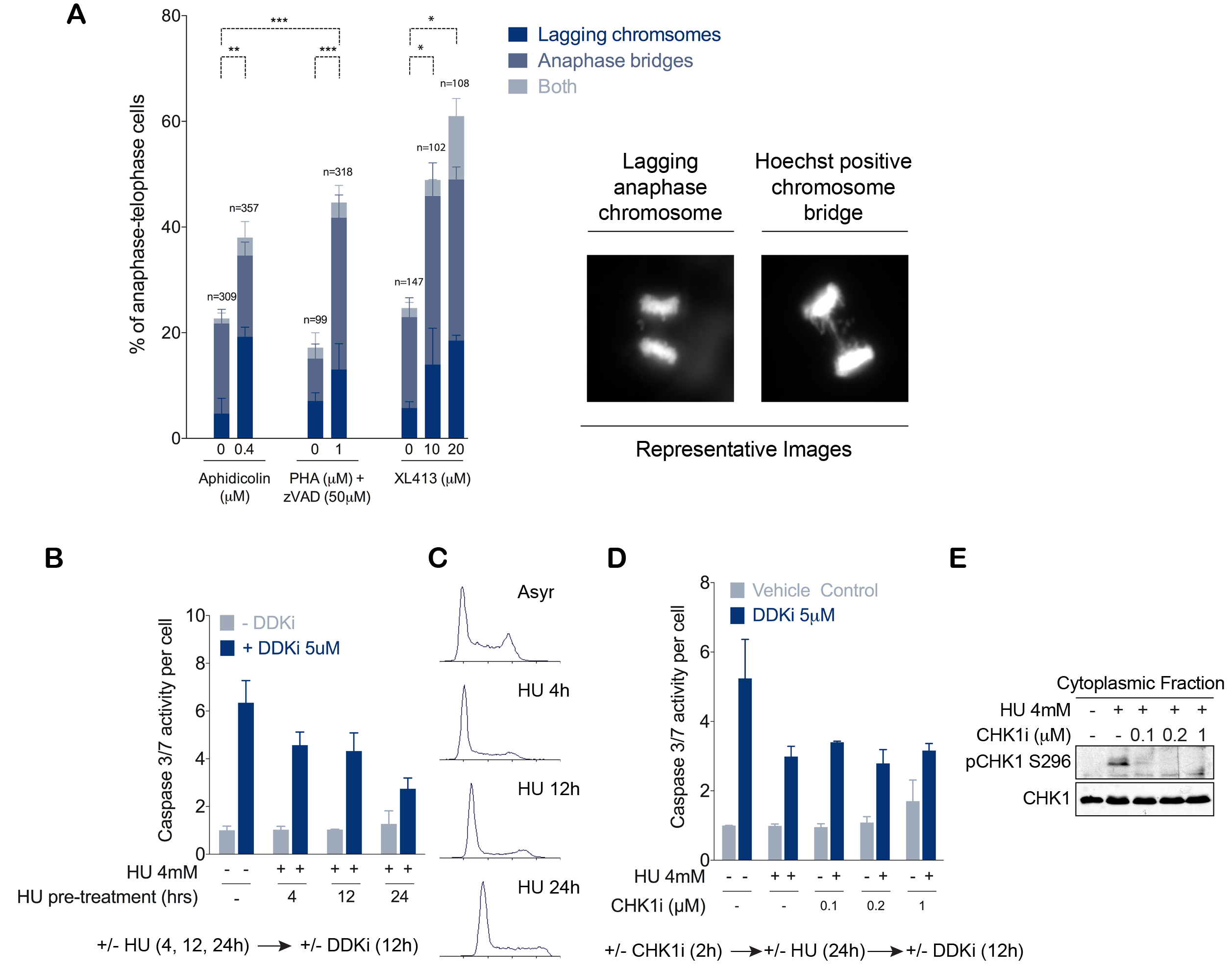
DDK inhibition induces mitotic abnormalities. **(A)** HCC1954 cells were treated with the indicated drugs for 24h and analyzed for mitotic abnormalities. Representative images are shown on the right and quantitated on the left. **(B, C)** HCC1954 cells were pre-treated with HU for the indicated times, analyzed by flow cytometry **(C)** or incubated with DDKi or DMSO for an additional 12h, and assayed for Caspase3/7 activity **(B)**. **(D, E)** HCC1954 cells were pre-treated with varying concentrations of CHK1i for 2h, exposed to HU for 24h, and then harvested for western blot **(E)** or treated with DDKi (dark bars) or DMSO (light bars) for an additional 12h prior to assaying for Caspase3/7 activity.

Fission yeast cells with hypomorphic DDK mutations also exhibited defective mitosis and DNA fragmentation [8, 10]. Importantly, these mutations are synthetically lethal with mutation in the cohesin protein *rad21* [8, 10]. These findings have led to the suggestion that DDK has a role in maintaining sister chromatid cohesion following DNA replication. The lack of checkpoint-induced cell cycle arrest, inability to restart naturally stalled forks, and compromised sister chromatid cohesion might explain the severe mitotic abnormalities seen in an asynchronous population of HCC1954 cells with reduced DDK activity.

In summary, we propose that ssDNA generated upon fork stalling is primarily a result of nascent strand degradation that requires DDK **(Figure 5)**. DDK is therefore required for the initiation of replication-checkpoint activation, and we also show for the recovery of stalled forks. An active replication-checkpoint would then attenuate nucleolytic activity at stalled forks to prevent excessive degradation of DNA by described mechanisms [41, 47–49]. Budding yeast cells lacking the checkpoint kinase Rad53, which inhibits DDK, exhibited extremely long tracks of ssDNA in presence of replication stress, which were abrogated upon deletion of Exo1 [33] again consistent with our model. Our analysis of EXO1 further shows that it plays a critical role in nascent strand degradation following exposure to HU and that DDK might regulate EXO1 stability and/or activity. The role of DDK in fork recovery would especially be important within origin poor regions of the genome where forks are known to stall. Since stalled forks cannot activate a robust checkpoint response in the absence of DDK, cells progress into M-phase with under-replicated DNA. Aberrant anaphase progression would result in chromosomal breakage, genomic instability, and might be the primary cause of cell death in DDKi-treated cancer cells **(Figure 5)**.

**Figure 5:**
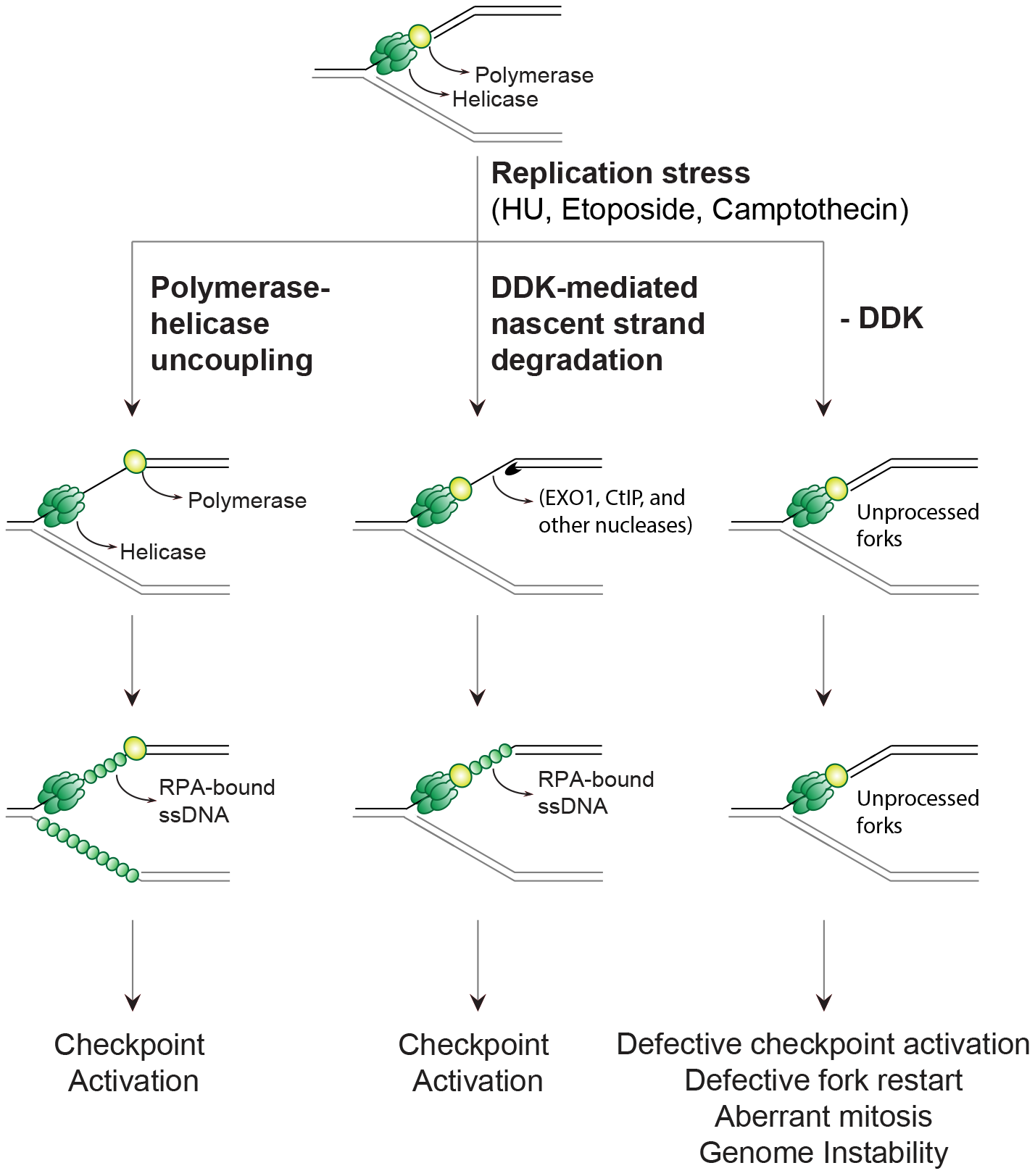
Model for role of DDK at stalled replication fork and events downstream of DDK inhibition. Replication stress results in stalled replication forks that accumulate RPA-bound ssDNA required for checkpoint activation. We propose DDK-mediated nucleolytic resection of newly synthesized DNA as a mechanism for generation of ssDNA at stalled replication forks. In the absence of DDK activity the forks remain unprocessed resulting in defects in checkpoint activation, fork restart, and cell cycle arrest.

## Author Contributions

N.K.S., F.C. and Y-L.L. performed all the experiments. J.P.M., P.P., and M.W. designed the experiments; N.K.S. and M.W. wrote the manuscript.

## Acknowledgments

We thank Bruce Stillman (Cold Spring Harbor) for the RPA antibodies. We thank the NIH National Cancer Institute grant R01CA197398 (to J.P.M.), the Van Andel Institute (M.W., N.K.S. and J.P.M.), and Michigan State University (N.K.S.) for support. Work in the Pasero laboratory is supported by the Institut National du Cancer (INCa) and the Ligue contre le Cancer (équipe labellisée).

---

**Supplementary Figure 1: DDK activity is required for CHK1 activation and RPA2 accumulation in response to various replication stresses. (A)** HCC1954 cells were pre-treated with DMSO or DDKi for the indicated times, then exposed to HU for 2h, followed by blotting for pCHK1 S317 or total CHK1. **(B, C)** HCC1954 cells were transfected with scrambled siRNA or CDC7 siRNA, treated with or without HU for 2h, harvested and subjected to cell fractionation followed by immunoblotting **(B)** or analyzed by immunofluorescence microscopy for RPA2 accumulation **(C). (D, E)** HCC1954 cells were pre-treated with DMSO or DDKi for 4h followed by incubation with or without camptothecin (CPT, **D**) or etoposide (Eto, **E**) for 2h and then subjected to cell fractionation and immunoblotting.

**Supplementary Figure 2: Replication-checkpoint pathway is not activated upon DDK inhibition. (A, B)** HCC1954 cells were treated with DMSO or DDKi for the indicated times and harvested for immunoblotting. Samples were fractionated for **(A)** and total cell extracts were used in **(B)**.

**Supplementary Figure 3: Role of DDK in the processing of stalled forks in response to replication stress. (A)** DNA fiber assay performed as in ‘Figure 2A’. Nascent strand resection was instead measured as the length of CldU tracks. **(B)** Experimental plan for DNA fiber assay in MCF7 cells. Cells were pre-treated with DMSO or DDKi for 1h, labelled consecutively with IdU and CldU (20 mins each), subjected to a thymidine chase with or without HU for 2h (still in the presence of DDKi or DMSO), and then harvested for DNA fiber assay. Nascent strand resection was measured either as CldU track length **(B)** or as a ratio of CldU to IdU incorporation **(C)**.

**Supplementary Figure 4: Depletion of EXO1, CtIP, and MRE11 does not affect the cell cycle profiles. (A, B)** HCC1954 cells were transfected with indicated siRNAs and 48 hours later treated with vehicle control (left) or HU (right) for 2h, and then harvested for western blot **(A)** or cell analysis by flow cytometry **(B)**.

**Supplementary Figure 5: EXO1 is required for HU-mediated nascent strand degradation. (A)** DNA fiber assay performed as in ‘Figure 3F’. Nascent strand resection was instead measured as the length of CldU tracks. **(B)** DNA fiber assay performed in the absence of thymidine chase. HU was added along with CldU and the length of IdU tracks were measured as an indicator of nascent strand degradation.

## Materials and Methods

### Cell lines and reagents

HCC1954 cells (ATCC) and Colo-205 (NCI-60) were cultured in RPMI-1640 media supplemented with 10% heat inactivated (HI) FBS, 50units/ml of penicillin, and 50μg/ml of streptomycin. The DDK inhibitors, PHA-767491 and XL413, were synthesized as described previously [18]. ATR inhibitor (VE-821, #A2521) and Camptothecin (#A2877) was from APExBIO. CHK1 inhibitor (LY2603618, #S2626) was from Selleckchem. Etoposide (#341205) was from EMD Millipore. Hydroxyurea (#H9120) was from USBiological. The antibodies were purchased as indicated: CST: PARP (#9542), pCHK1 S317 (#12302), pCHK1 S345 (#2348), pCHK1 S296 (#2349), CHK1 (#2360), pCHK2 T68 (#2197), CHK2 (#6334), RAD51 (#8875), CtIP (#93110), RECQL4 (#2814); Bethyl Laboratories Inc.: pMCM2 S53 (A300-756A), pMCM2 S108 (A300-094A), MCM2 (A300-122A), pRPA2 S33 (A300-246A), pRPA2 S4/S8 (A300-245A), ORC2 (A302-735A), EXO1 (A302-639A), MRE11 (A303-998A), BLM (A300-110A), RECQL5 (A302-520A), RIF1 (A300-568A); Santa Cruz Biotechnology: WRN (sc-376182), RECQL1 (sc-166388); MBL International Corporation: CDC7 (K0070-3S); Sigma: β-actin (A5441), Tubulin (T9026); antibodies against RPA1 (NA13, EMD Millipore) and RPA2 (04-1481, EMD Millipore) were gifts from Dr. Bruce Stillman; GE Healthcare: anti-mouse-HRP (NA931V), and anti-rabbit-HRP (NA934V).

### RNAi interference

HCC1954 cells were plated in 6-well plates (75000 cells/well) allowed to grow for 36h before transfection. siRNA transfection was performed with Lipofectamine RNAiMAX (Invitrogen) according to manufacturer’s instructions. Each well was transfected with 2μl transfection reagent and a final siRNA concentration of 25nM (*CDC7*, *EXO1*) or 5nM (*MRE11*, *CtIP*, *BLM, RECQL1, RECQL4, RECQL5, WRN, RIF1*) in a total volume of 2ml. Media was replaced 24 hours after transfection and the cells were either harvested or exposed to indicated treatments 48h after transfection. Following siRNAs were used: *CDC7* (CDC7-L1, Dharmacon custom siRNA, GGCAAGATAATGTCATGGGA), *EXO1* (Qiagen, SI02665145, GAUGUAGCACGUAAUUCAAtt), *MRE11A* (Thermo Scientific, #s8960, CCCGAAAUGUCACUACUAAtt), *CtIP* (Thermo Scientific, #s142451, CGAAUCUUAGAUGCACAAAtt), *BLM* (Thermo Scientific #s1999, GAUAUCUUCCAAAACGAAAtt).

### Immunoblotting and protein fractionation

Whole cell extracts were prepared by re-suspending the pellets in RIPA buffer (150mM NaCl, 1% NP-40, 0.5% sodium deoxycholate, 0.1% SDS, 50mM Tris-HCl, pH 8) containing protease inhibitors (100μM PMSF, 1mM Benzamidine, 2.5μg/ml Pepstatin A, 10μg/ml Leupeptin, and 10μg/ml Aprotinin) and phosphatase inhibitors (1mM each NaF, Na_3_VO_4_, Na_2_P_2_O_7_). Protein concentration was measured using the BCA protein assay kit (Pierce, #23227). Cell fractionation into cytosolic, nuclear soluble, and nuclear insoluble (chromatin) fractions was performed as described previously [50]. Pellets were re-suspended in lysis Buffer A (10mM HEPES (pH 7.9), 10mM KCl, 1.5 mM MgCl_2_, 0.34M Sucrose, 10% Glycerol, 1mM DTT, and protease and phosphatase inhibitors) and Triton X-100 was added to a final concentration of 0.1%. After incubation on ice for 8min, lysates were centrifuged at 1,300g, at 4⁰C, for 5min. The supernatant was collected and clarified by high-speed centrifugation (20,000g, 4⁰C, 5min) to obtain cytosolic fraction. The pellet was washed once with Buffer A and then lysed in Buffer B (3mM EDTA, 0.2 mM EGTA, 1mM DTT, protease and phosphatase inhibitors) for 30min on ice. Soluble nuclear fraction (supernatant) was collected by centrifugation at 1,700g, at 4⁰C, for 5min. The chromatin fraction (pellet) was washed once with Buffer B, re-suspended in Buffer B, and sonicated briefly. Protein concentration in each fraction was measured using Bradford assay (Bio Rad, #500-0006). Equal amounts of proteins were subjected to SDS-PAGE and transferred to nitrocellulose membrane (Millipore, HATF304F0). Transfer efficiency and equal loading was confirmed by Ponceau S staining. Membranes were blocked overnight at 4⁰C with 5% non-fat milk in TBS-T, followed by incubation in primary and secondary antibodies (1h at RT, 2% milk in TBS-T). Protein bands were visualized using SuperSignal West Pico solutions (Thermo Scientific).

### Analysis of Caspase 3/7 activity

5000 cells per well were plated in 96 well plates. 24 hours later cells were treated and incubated for the indicated period of time at 37⁰C. Caspase 3/7 activity and viable cell number were then measured using the Caspase-Glo 3/7 assay (Promega) and CellTiter-Glo assay (Promega), respectively. The ‘caspase activity per cell’ was obtained by normalizing total caspase activity to cell number. Luminescence was measured using BioTek Synergy Microplate Reader 30 minutes after addition of ‘Glo’ reagents.

### Cell Cycle Analysis

Cells were trypsinized, washed twice with cold PBS, and fixed/permeabilized in 70% ice-cold ethanol (made in water). After fixation on ice for 30mins cells were centrifuged at 400g (4⁰C, for 5mins), washed once with cold PBS, and centrifuged again. The pellets were re-suspended in analysis buffer (10μg/ml propidium iodide and 250μg/ml RNAase) and incubated at 37⁰C for 30min. Cell cycle profiles were obtained using FACSCalibur^TM^ (BD Biosciences) flow cytometer. The data was analyzed using Flowing Software.

### Immunofluorescence

HCC1954 cells were seeded at 50,000 to 70,000 cells per well on number 1.5 coverglass in 24-well tissue culture dishes. siRNA-mediated knockdown was carried out in 6-well plates. 24 hours later the cells were trypsinized and re-plated on coverglass in 24-well plates. After 36 hours, cells were treated as indicated. For RPA2 and BrdU immunofluorescence analysis, cells were washed with 1X PBS, pre-extracted with 0.2% Triton X-100 (in PBS) for 5min, washed twice with 1X PBS, and fixed in 4% paraformaldehyde for 10 minutes at room temperature. Cells were then stained with primary antibody (1:1000 in DMEM + 10%FBS + 0.01% sodium azide) overnight at 4°C on a shaker. After 3 5-minute washes with PBS-T (0.01% Tween 20) cells were incubated in Alexa Fluor 488 secondary antibody (# A-11001, 1:1000 in DMEM + 10%FBS + 0.01% sodium azide) for 1h followed by 3 washes with PBS-T. Nuclear DNA was stained with Hoechst 33342 (2 μg/ml). Coverglass was inverted onto microscope slides using mounting gel. Cells were imaged using a 60X oil-immersion objective on a Nikon Eclipse Ti florescent microscope. Image processing and quantification were completed with NIS Elements software (Nikon). To quantify, regions of interest (ROI) were drawn around each nucleus. Using HU-treated cells as a positive control, intensity thresholds (binary) were set to include all pixels equal to or greater than the intensity of the background 488 fluorescence. The binary was applied to all samples and the sum intensity of the ‘binary in ROI’ was calculated for each nucleus. All values were normalized to the mean intensity of the vehicle control. For analysis of mitotic abnormalities, cells were fixed and permeabilized in 70% ethanol. Nuclear DNA was stained with Hoechst 33342 (2ug/ml). Image analysis was performed using NIS Elements software (Nikon). Statistical analysis was performed using Graph Pad Prism.

### DNA fiber spreading

DNA fiber spreading was performed as described previously [51, 52]. Briefly, sub-confluent cells were sequentially labeled first with 10 μM 5-iodo-2’-deoxyuridine (IdU) and then with 100 μM 5-chloro-2’-deoxyuridine (CldU) for the indicated times. One thousand cells were loaded onto a glass slide (Star Frost) and lysed with spreading buffer (200 mM Tris-HCl pH 7.5, 50 mM EDTA, 0.5% SDS) by gently stirring with a pipette tip. The slides were tilted slightly and the surface tension of the drops was disrupted with a pipette tip. The drops were allowed to run down the slides slowly, then air-dried, fixed in methanol/acetic acid 3:1 for 10 minutes, and allowed to dry. Glass slides were processed for immunostaining with mouse anti-BrdU to detect IdU (347580, BD Biosciences), rat anti-BrdU (ABC117-7513, Eurobio Abcys) to detect CldU, mouse anti-ssDNA (MAB3868, Millipore) antibodies and corresponding secondary antibodies conjugated to various Alexa Fluor dyes. Nascent DNA fibers were visualized by using immunofluorescence microscopy (Zeiss Apotome 2). The acquired DNA fiber images were analyzed by using MetaMorph Microscopy Automation and Image Analysis Software (Molecular Devices) and statistical analysis was performed with Graph Pad Prism (Graph Pad Software). The lengths of at least 150 IdU/CldU tracks were measured per sample.

